# Continuous Multiplexed Phage Genome Editing Using Recombitrons

**DOI:** 10.1101/2023.03.24.534024

**Authors:** Chloe B. Fishman, Kate D. Crawford, Santi Bhattarai-Kline, Karen Zhang, Alejandro González-Delgado, Seth L. Shipman

## Abstract

Bacteriophages, which naturally shape bacterial communities, can be co-opted as a biological technology to help eliminate pathogenic bacteria from our bodies and food supply^1^. Phage genome editing is a critical tool to engineer more effective phage technologies. However, editing phage genomes has traditionally been a low efficiency process that requires laborious screening, counter selection, or *in vitro* construction of modified genomes^2^. These requirements impose limitations on the type and throughput of phage modifications, which in turn limit our knowledge and potential for innovation. Here, we present a scalable approach for engineering phage genomes using recombitrons: modified bacterial retrons^3^ that generate recombineering donor DNA paired with single stranded binding and annealing proteins to integrate those donors into phage genomes. This system can efficiently create genome modifications in multiple phages without the need for counterselection. Moreover, the process is continuous, with edits accumulating in the phage genome the longer the phage is cultured with the host, and multiplexable, with different editing hosts contributing distinct mutations along the genome of a phage in a mixed culture. In lambda phage, as an example, recombitrons yield single-base substitutions at up to 99% efficiency and up to 5 distinct mutations installed on a single phage genome, all without counterselection and only a few hours of hands-on time.

## INTRODUCTION

Bacteriophages (phages) naturally control the composition of microbial ecosystems through selective infection of bacterial species. Humans have long sought to harness this power of phages to make targeted interventions to the microbial world, such as delivering phages to a patient suffering from an infection to eliminate a bacterial pathogen. This approach to mitigate pathogenic bacterial infections, known as phage therapy, predates the discovery of penicillin, with 100 years of evidence for efficacy and safety^4^. However, the success of small molecule antibiotics over the same period of time has overshadowed and blunted innovation in phage therapy.

Unfortunately, it is now clear that reliance on small molecule antibiotics is not a permanent solution. Antimicrobial resistance in bacteria was associated with between 1–5 million deaths worldwide in 2019^5^, a figure that is projected to rise in the coming decades^6^. As such, there is a pressing need for alternatives or adjuvants to small-molecule antibiotics, like phage therapy, to avoid returning to the rampant morbidity of bacterial infections of the pre-antibiotic era. This work has already begun, with researchers and clinicians utilizing phage screening pipelines to identify natural phages for use in patients to overcome antimicrobial-resistant infections^7,8^.

While these efforts demonstrate the potential of phage therapeutics, they do not scale well. Screening natural phages for each patient is time- and effort-intensive, requires massive repositories of natural phages, and results in the clinical use of biological materials that are not fully characterized^1^. To functionally replace or supplement small-molecule antibiotics, phage therapy needs to be capable of industrialization and more rapid iteration. This will likely include modifying known phages to create engineered therapeutic tools that target specific pathogens and evade natural bacterial immunity, rather than just the opportunistic isolation of natural phages. However, such approaches are limited by the relative difficulty in modifying phage genomes^1^.

The various approaches to modify phage genomes and the limitations imposed by each have been recently reviewed^2^. One approach is to modify phage genomes by recombination within their bacterial host, which is inefficient and requires laborious screening of phage plaques. That screening effort can be reduced by imposing a counterselection on the unedited phage, such as CRISPR-based depletion of the wild-type phage^9–12^. However, this negative selection is not universally applicable to all edit types because it requires functional disruption of a protospacer sequence^13^. Phages also frequently escape CRISPR targeting^14^, which can result in most selected phages containing escape mutations outside of the intended edit^9^. Another approach is to “reboot” a phage by assembling a modified phage genome *in vitro* and repackaging it in a host^15^, but such rebooting requires extensive work to enable *in vitro* assembly of a phage genome, and phage genome size limits the efficiency of transformation. A fully cell-free packaging system eliminates the issue of inefficient transformation^16^, but instead requires substantial upfront technical development, which is host species-specific.

Because of the urgent need for innovation in phage therapeutics and the clear technical hurdle in modifying phage genomes, we developed an alternative system for phage engineering. Our system edits phage genomes as they replicate within their bacterial hosts, by integrating a single-stranded DNA (ssDNA) donor encoding the edit into the replicating phage genome using a single-stranded annealing protein (SSAP) and single-stranded binding protein (SSB). A critical innovation is that we use a modified bacterial retron^17–20^ to continuously produce the editing ssDNA donor by reverse transcription within the host. Thus, propagating a population of phage through this host strain enables continuous accumulation of the intended edit over generations within a single culture.

Moreover, this system enables a more complex form of editing in which the bacterial culture is composed of multiple, distinct editing hosts, each producing donors that edit different parts of the phage genome. Propagation of phages through such a complex culture leads to the accumulation of multiple distinct edits at distant locations in individual phage genomes. Here, we demonstrate this approach, showing that (1) the editing is a continuous process in which edits accumulate over time; (2) it can be applied to multiple types of phage and used to introduce different edit types; (3) it can be optimized to reach efficiencies that do not require counterselection; and (4) it can be used to make multiplexed edits across a phage genome. For disambiguation with other techniques, we call this approach *phage retron recombineering*, and term the molecular components that include a modified retron a *recombitron*.

## RESULTS

### Recombitrons Target Phage Genomes for Editing

There are four core molecular components of a recombitron: a retron non-coding RNA (ncRNA) that is modified to encode an editing donor; a retron reverse transcriptase (RT); a single-stranded binding protein (SSB); and a single-stranded annealing protein (SSAP) (**Fig 1a**). Endogenous retrons partially reverse transcribe a short (∼200 base) ncRNA into a single-stranded reverse-transcribed DNA (RT-DNA) of ∼90 bases. In bacteria, this RT-DNA is used in conjunction with retron accessory proteins to detect and respond to phage infection. The retron accessory proteins are necessary for the phage defense phenotype and are not included in the recombitron to avoid reconstituting an anti-phage system^21–24^. We modify the retron ncRNA by adding nucleotides to the region that is reverse transcribed that are homologous to a locus in the phage genome and carry the edit we aim to incorporate.

**Figure 1.**
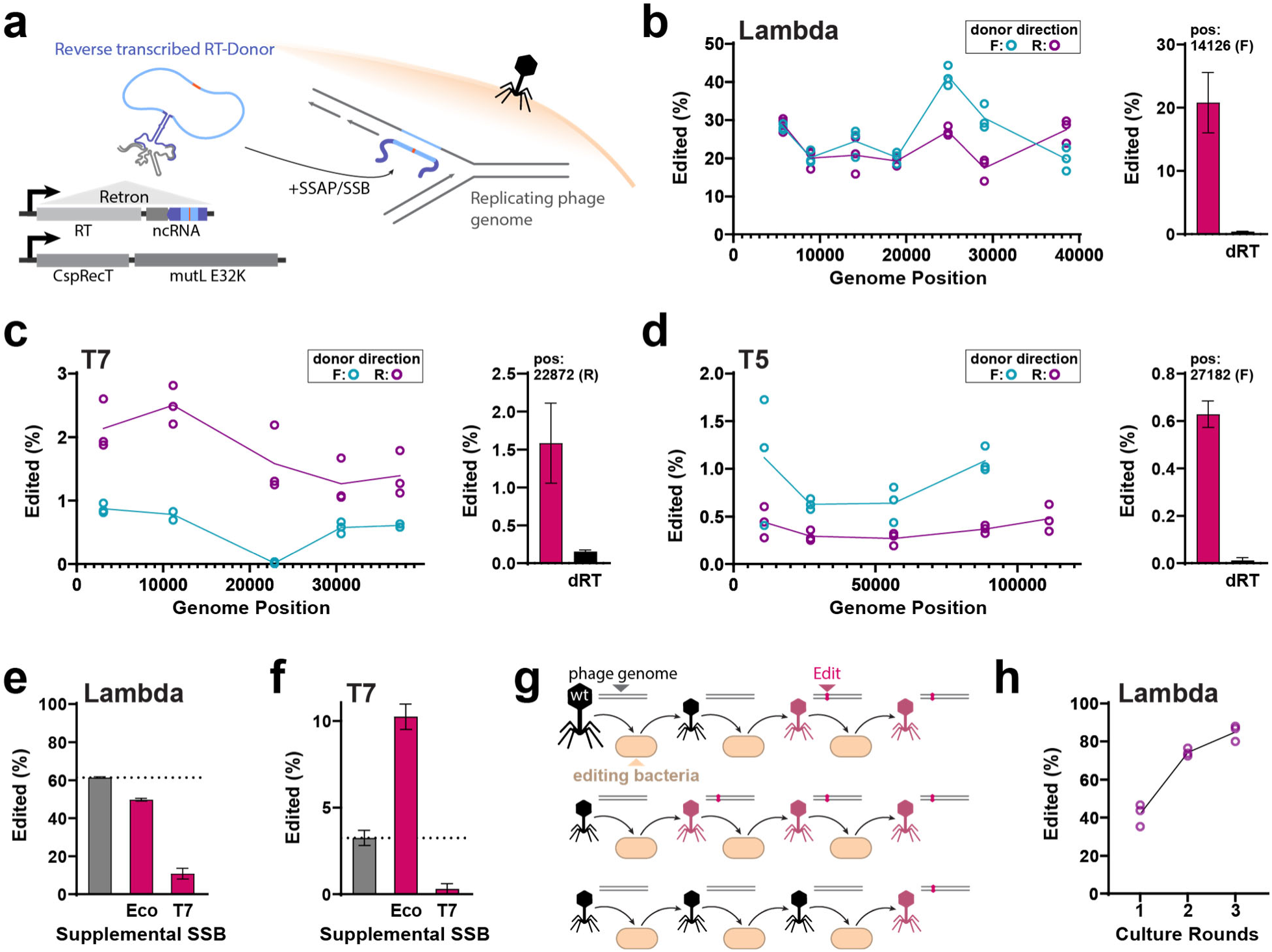
Recombitrons Target Phage Genomes for Continuous Editing. **a.** Recombitron schematic: a modified retron generates ssDNA containing a donor sequence with an edit flanked by homology to the phage genome that is integrated into the phage genome during replication by SSB and SSAP. The retron cassette is expressed from an operon containing a reverse transcriptase (RT) and ncRNA. The reverse-transcribed region of the ncRNA is shown in purple with an inserted donor sequence in light blue, and the edit site is shown in orange. A second operon expresses CspRecT and mutL E32K. **b.** Left: Edited phage genomes (as a % of all genomes) across lambda phage. Editing with a forward RT-DNA is shown in blue, reverse in purple. Three individual replicates are shown in open circles at each point. Right: Editing with the recombitron at site 14,126 (±SD) is significantly greater than editing with a dRT control (t-test, *P*=0.0018). **c.** Left: Edited phage genomes across phage T7 for three replicates at each position, displayed as in b. Right: Editing with the recombitron at site 22,872 (±SD) is significantly greater than editing with a dRT control (t-test, *P*=0.0094). **d.** Left: Edited phage genomes across phage T5 for three replicates at each position, displayed as in b. Right: Editing with the recombitron at site 27,182 (±SD) is significantly greater than editing with a dRT control (t-test, *P*<0.0001). **e.** Editing of lambda at site 30,840 (F) (±SD) compared to editing with supplemental expression of *E. coli* SSB or T7 SSB. There is a significant effect of SSB expression (one-way ANOVA, *P*<0.0001, N=3), with both the *E. coli* (*P*=0.005) and T7 (*P*=<0.0001) significantly different than the no SSB condition (Dunnett’s, corrected). **f.** Editing of T7 at site 11,160 (R) (±SD) compared to editing with supplemental expression of *E. coli* SSB or T7 SSB. There is a significant effect of SSB expression (one-way ANOVA, *P*<0.0001, N=3), with both the *E. coli* (*P*=0.0002) and T7 (*P*=0.0127) significantly different than the no SSB condition (Dunnett’s, corrected). **g.** Schematic illustrating the accumulation of edited phages with multiple rounds of editing. **h.** Proportion of edited lambda phage increases over 3 rounds of editing of lambda at site 30,840 (F). Three replicates per round are shown in open circles. Additional statistical details in Supplementary Table 1.

The recombitron we begin with here specifically contains a modified retron-Eco1 ncRNA expressed on the same transcript as a retron-Eco1 RT, which will produce a 90-base-long reverse-transcribed editing donor (RT-Donor) inside the host bacteria. Once produced, the SSB will bind the editing RT-Donor to destabilize internal helices and promote interaction with an SSAP. In this case, we leverage the endogenous *E. coli* SSB. Next, an SSAP promotes annealing of the RT-Donor to the lagging strand of a replication fork, where the sequence is incorporated into the newly replicated genome^25^. From a separate promoter, we express the SSAP CspRecT along with an optional recombitron element mutL E32K, a dominant-negative version of *E. coli* mutL that suppresses mismatch-repair^26–28^. Such suppression is necessary when creating single-base mutations, but not required for larger insertions or deletions.

This system benefits from stacking several beneficial modifications to the retron and recombineering machinery discovered previously. The retron ncRNA, in addition to being modified to encode an RT-Donor, is also modified to extend the length of its a1/a2 region, which we previously found to increase the amount of RT-DNA produced^20,29^. The SSAP, CspRecT, is more efficient than the previous standard, lambda β, and is known to be compatible with *E. coli* SSB^27^. The dominant-negative mutL E32K eliminates the requirement to pre-engineer the host strain to remove mutS^26,30^. A previous study attempted a single edit using a retron-produced donor against T5 phage, but only reported editing after counterselection^12^. By using these stacked innovations, we aim to produce a system that does not require counter selection.

To test whether recombitrons can be used to edit phages, we designed recombitrons targeting 5-7 sites across the genome of four *E. coli* phages: Lambda, T7, T5, and T2. Each recombitron was designed to make synonymous single-base substitutions to a stop codon, which we assume to be fitness neutral. We constructed separate recombitrons to produce the RT-Donor in either of the two possible orientations, given that the mechanism of retron recombineering requires integration into the lagging strand, and created a recombitron with a catalytically-dead RT at one position per phage as a control. We pre-expressed the recombitron for 2 hours in BL21-AI *E. coli* that lack the endogenous retron-Eco1, then added phage at an MOI of 0.1 and grew the cultures for 16 hours overnight. The next day, we spun down the cultures to collect phage in the supernatant. We used PCR to amplify the regions of interest in the phage genome (∼300 bases) surrounding the edit sites. We sequenced these amplicons on an Illumina MiSeq and quantified editing with custom software.

We observed RT-dependent, counter-selection-free editing in lambda (∼20%), T7 (∼1.5%), and T5 (∼0.6%) (**Fig 1b-d**). We did not observe editing in T2 above the dead RT background level, which could be due to T2’s use of modified 5-hydroxymethycytosines (**Supplementary Fig 1a**)^31^. In lambda, T7, and T5, we obtained similar maximum editing efficiencies at each site across the genome. Editing efficiency was affected by RT-Donor direction in a manner generally consistent with individual phage replication mechanisms. Lambda has an early bidirectional phase of replication and a later unidirectional phase^32^. Thus, either strand may be lagging at some point and, accordingly, we observed similar efficiencies for the forward and reverse recombitron (**Fig 1b**). T7 has a single origin of replication at one end of its linear genome, and we observed directional editing favoring the predicted lagging (reverse) strand (**Fig 1c**)^33^. Interestingly, T5 replication is less well studied, with one report describing bidirectional replication from multiple origins^34^. However, we found a clear strand preference indicating dominant unidirectional replication from one end of the phage genome (**Fig 1d**).

Editing efficiency differed among the phages tested, with lambda being the most efficiently edited. One possible explanation for this difference is that lambda is the only temperate phage tested, so hypothetically the editing could be occurring while lambda was integrated into the *E. coli* genome. However, we obtained two results inconsistent with this hypothesis. First, we generated a lambda strain with an inactivated cI gene (ΔcI) that is required for prophage maintenance, and found similar levels of editing in that lambda strain as compared with our standard lambda strain (**Supplementary Fig 1b**)^35^. Next, we modified the bacterial host to remove the lambda prophage integration site (attB), and again found similarly high levels of editing of the ΔcI strain in the attB^-^ bacteria (**Supplementary Fig 1b**)^36^. Another possibility is that lambda β, an SSAP encoded by lambda, assists the recombitron^25^. However, we found no editing of lambda in the absence of CspRecT expression, which is inconsistent with that possibility (**Supplementary Fig 1c**). Perhaps the recombitron differentially affects phage replication, which in turn affects editing efficiency. However, we found no effect on replication of any phage (as measured by phage titer) when the recombitron was included or expressed in the phage’s bacterial host (**Supplementary Fig 1d,e**).

Another alternative explanation is SSB compatibility^37^. Phage T7, unlike lambda, encodes a separate SSB in its genome (gp2.5)^38^. Perhaps T7 SSB competes with the *E. coli* SSB for the recombitron RT-DNA, which could inhibit interactions with the CspRecT. To test this possibility, we repeated the lambda and T7 editing experiments at one locus each, while overexpressing either the *E. coli* or the T7 SSB. Consistent with this explanation, we found that overexpression of the T7 SSB has a large negative impact on editing of lambda, while the *E. coli* SSB had a much smaller impact (**Fig 1e**). We similarly found that overexpression of the T7 SSB strongly reduced editing of T7, whereas overexpression of the E. coli SSB had a large positive effect on T7 editing, more than doubling the efficiency to a rate of 10% (**Fig 1f**). Thus, the T7 SSB appears to inhibit the retron recombineering approach, but can be counteracted by overexpressing a compatible SSB. We tried a similar approach with T5, which has a much less characterized SSB (PC4-like, by homology to phage T4^39^). While the T7 SSB similarly inhibited T5 editing, the T5 PC4-like protein had a small negative effect on editing, and the *E. coli* SSB did not improve T5 editing (**Supplementary Fig 1f-h**).

### Continuous Editing

One clear advantage of using recombitrons for phage editing is the fact that they edit continuously while the phage is propagating through the culture, increasing the proportion of edited phages with every generation (**Fig 1g**). This is quite distinct from other methods of recombineering in phages where materials are delivered via transformation of the host at a single point in time^40^. To illustrate the continuous nature of this approach, we edited lambda over three rounds of culture. Editing and analysis was performed as described above, but at the end of each round of editing, we propagated the resulting phage population through a fresh culture of editing the host. The percentage of edited genomes increased with each additional round, reaching >80% of phages edited after three rounds (**Fig 1h**).

### Optimizing Recombitron Parameters

Given the initial success of recombitrons, we next tested the parameters of the system to achieve optimal editing. The first parameter we tested was length of the RT-Donor. We designed a set of recombitrons with different RT-Donor lengths, each encoding a lambda edit (C14070T) in the center of the homologous donor (**Fig 2a**). We found no editing with a 30 base RT-Donor, but all other lengths tested from 50 to 150 bases produced successful editing (**Fig 2b**). The highest overall editing rates occurred with a 70 base RT-Donor. However, for donors between 50 and 150 bases, an analysis corrected for multiple comparisons only found a significant difference between a 70 and 150 base donors, indicating that long RT-Donor length is detrimental to editing efficiency (**Fig 2b**).

**Figure 2.**
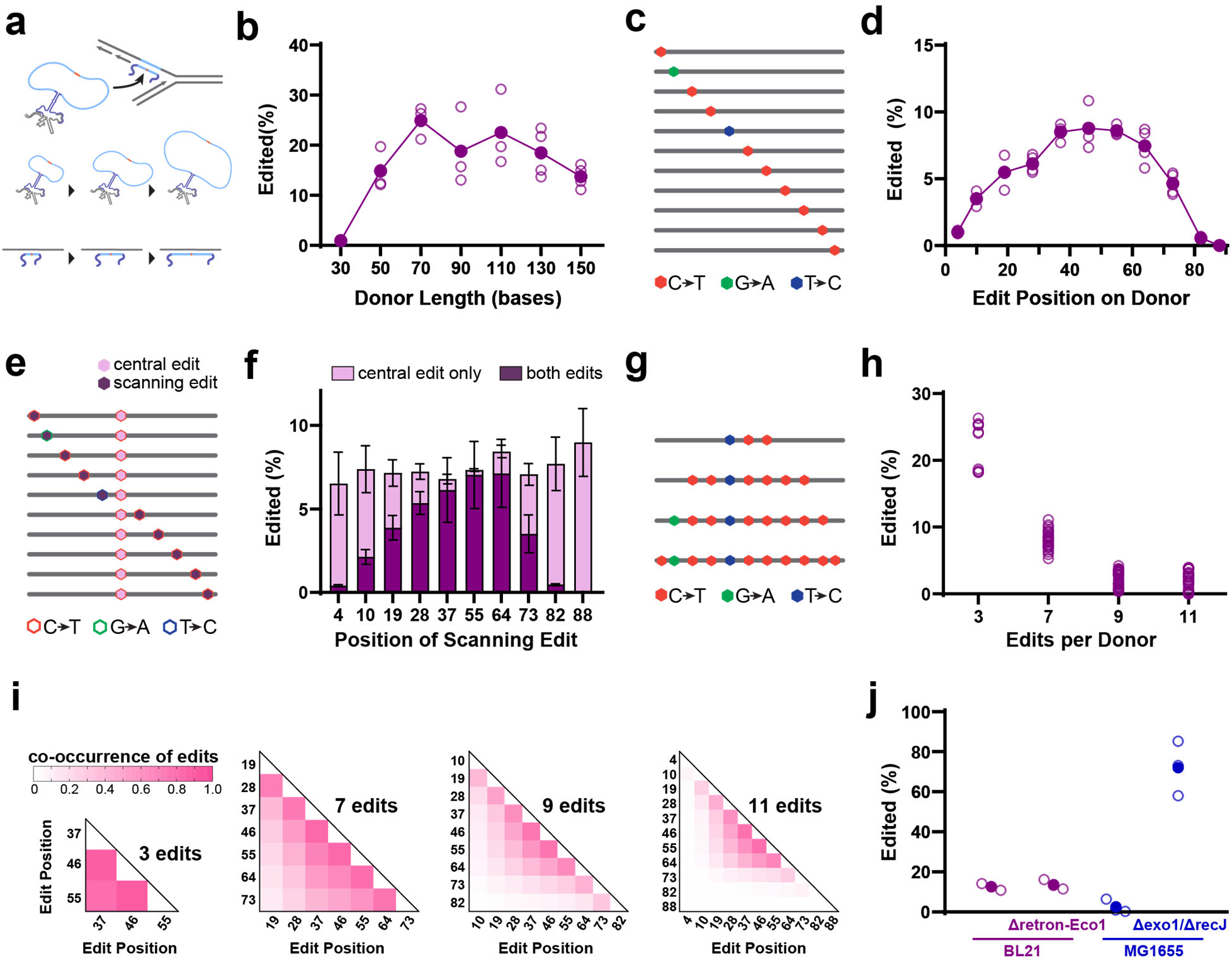
Optimizing Recombitron Parameters with Lambda Phage. **a.** Schematic of recombitron with longer and shorter RT-Donors, with more or less homology to the genome surrounding the edit site. **b.** Editing (%) of lambda with donors ranging in size from 30-150 bases. Three to four biological replicates are shown in open circles and closed circles show the average per recombitron, demonstrating an effect of length (one-way ANOVA, *P*<0.0001). All RT-Donors greater than 50 bases yield significantly improved editing from a 30 base donor (Sidak’s corrected, *P*<0.05), and a RT-Donor of 150 bases is significantly worse than 70 bases (Sidak’s corrected, *P*<0.05). **c.** Schematic of RT-Donors with homology to a single lambda region, each containing a scanning edit. **d.** Editing (%) of lambda with donors containing scanning edit. Four biological replicates are shown in open circles and closed circles show the average per recombitron, demonstrating an effect of edit placement (one-way ANOVA, *P*<0.0001). Edit placement at positions ≤28 or ≥73 is significantly worse position 46 (Dunnett’s corrected, *P*<0.001). **e.** Schematic of RT-Donors containing the scanning edit with an additional, constant central edit. **f.** Editing (%) of lambda with donors containing both the scanning and central edits. Editing rate when only the central edit is acquired is shown in light pink, acquisition of both edits shown in purple (±SD). Position of scanning edit does not affect total editing (one-way ANOVA, *P*=0.7868), but does affect whether both edits are installed (one-way ANOVA, *P*<0.0001). **g.** Schematic of multi-edit recombitrons containing 3, 7, 9, and 11 edits per RT-Donor. **h.** Editing (%) of lambda from the multi-edit recombitrons, each site across three to four biological replicates is shown as an open circle. Number of edits per RT-Donor significantly affects average editing rate across sites (one-way ANOVA, *P*<0.0001), with 7, 9, and 11 edits all performing significantly worse than 3 (Dunnett’s corrected, *P*<0.0001). **i.** Co-occurrence of edits that are acquired from multi-edit donors. Heatmap shows correlation coefficient r^2, normalized to the overall editing rate for each donor at each site. **j.** Editing (%) of lambda in different host strains at position 14,126 (R), using a plac-recombitron that is functional across all strains. Three biological replicates are shown as open circles and the average is shown as a closed circle. There is a significant effect of strain (one-way ANOVA, *P*<0.0001). The MG1655 (exo1-/recJ-) strain yielded significantly more editing than the BL21 (Dunnett’s corrected, *P*<0.0001). Additional statistical details in Supplementary Table 1.

The next parameter we tested was positioning of the edit within the RT-Donor. We tested a set of recombitrons where we held a 90 base region of homology to lambda constant and encoded an edit at different positions along the donor, each of which introduce a distinct synonymous single-base substitution (**Fig 2c**). Here we found that the editing rate increased as the edit approached the center of the donor (**Fig 2d**). After correcting for multiple comparisons, we found no difference between edit placement between a central 27 bases from position 37 to 64, but a significant decline in editing outside the central region.

We also tested the effect of position when encoding multiple edits on a single RT-Donor. In this case, we tested a set of recombitrons with edits at different positions along the RT-Donor as above, but additionally included a central edit on every RT-Donor (**Fig 2e**). Similar to the effect above, we found that the rate of incorporating both of the edits on a single phage genome declined as the scanning edit approached either edge of the RT-Donor (**Fig 2f**). However, there was a consistent editing rate for all recombitrons in phage genomes that were only edited at the central site. When both edits were positioned toward the center of the donor, most genomes contained both edits, whereas when the scanning edit was positioned toward the edge of the RT-Donor, there was a significant increase in likelihood of incorporating only the central edit (**Fig 2f**). We interpret this result as evidence that the RT-Donor can be partially used, with a bias toward using the central part of the RT-Donor. We also found only a very low rate of the scanning edit being incorporated without the central edit (**Supplementary Fig 2**).

Next, we tested the effect of increasing the number of edits per recombitron. We constructed a set of recombitrons with 3, 7, 9, or 11 of the scanning edits used above (**Fig 2g**). We found that editing with the 3-edit recombitron was comparable to the single edits we previously tested, but editing rates declined across all edit sites when additional edits were encoded on a single recombitron (**Fig 2h**). We interpret this as an effect of decreasing homology across the donor, making it more difficult to anneal to the target site. To further assess which edits on the same RT-Donor were likely to be acquired together, we analyzed the co-occurrence of edits. Because of the effect of decreasing homology on the editing rate, we calculated the correlation coefficient r^2 and normalized it to the overall editing rate for each donor at each site. We found that there was a bias toward acquiring the more central edits overall, and a tendency to co-edit nearby sites (**Fig 2i**), again supporting the potential for partial use of the RT-Donor.

### Optimizing host strain

We also tested the effect of the editing host. Specifically, we compared editing in: B-strain *E. coli* (BL21-AI); a derivative of BL21-AI lacking the endogenous retron-Eco1; K-strain *E. coli* (MG1655); and derivative of MG1655 lacking Exo1 and RecJ – nucleases whose removal was previously shown to increase recombination rates using synthetic oligonucleotides^18,41^. We found no difference between the BL21-AI and retron-deletion derivative, indicating that the endogenous retron does not interfere with the recombitron system (**Fig 2j**). We found decreased editing in the wild-type K-strain that was not statistically significant as compared to the B-strain, but significantly improved editing in the modified K-strain versus the B-strain (**Fig 2j**).

### Insertions and Deletions via Recombitrons

Genomic deletions and insertions are also useful for engineered phage applications. Deletions can be used to remove potential virulence factors or eukaryotic toxins from phage genomes or to optimize phages by minimization. Insertions can be used to deliver cargo, such as nucleases that can help kill target cells, or anti-CRISPR proteins to escape phage defense systems. Therefore, we tested the efficiency of engineering deletions and insertions of increasing size into the lambda genome.

We began by comparing the efficiency of deleting 2, 4, 8, 16, or 32 bases to one of the single base synonymous substitutions we tested previously (**Fig 3a**). These experiments were performed in the engineered K-strain that we found to yield higher editing efficiencies using inactivated cI lambda as the phage. We deleted bases preceding position 37,673 using recombitrons where a 90 base RT-Donor contained flanking homology around the deletion site, but omitted the bases to be deleted. We found consistent editing of ∼45% for each of the deletions (**Fig 3b**) using Illumina amplicon sequencing. The deletion size did not significantly affect the efficiency of editing, although all the deletions exhibited significantly lower editing than the synonymous single base substitution.

**Figure 3.**
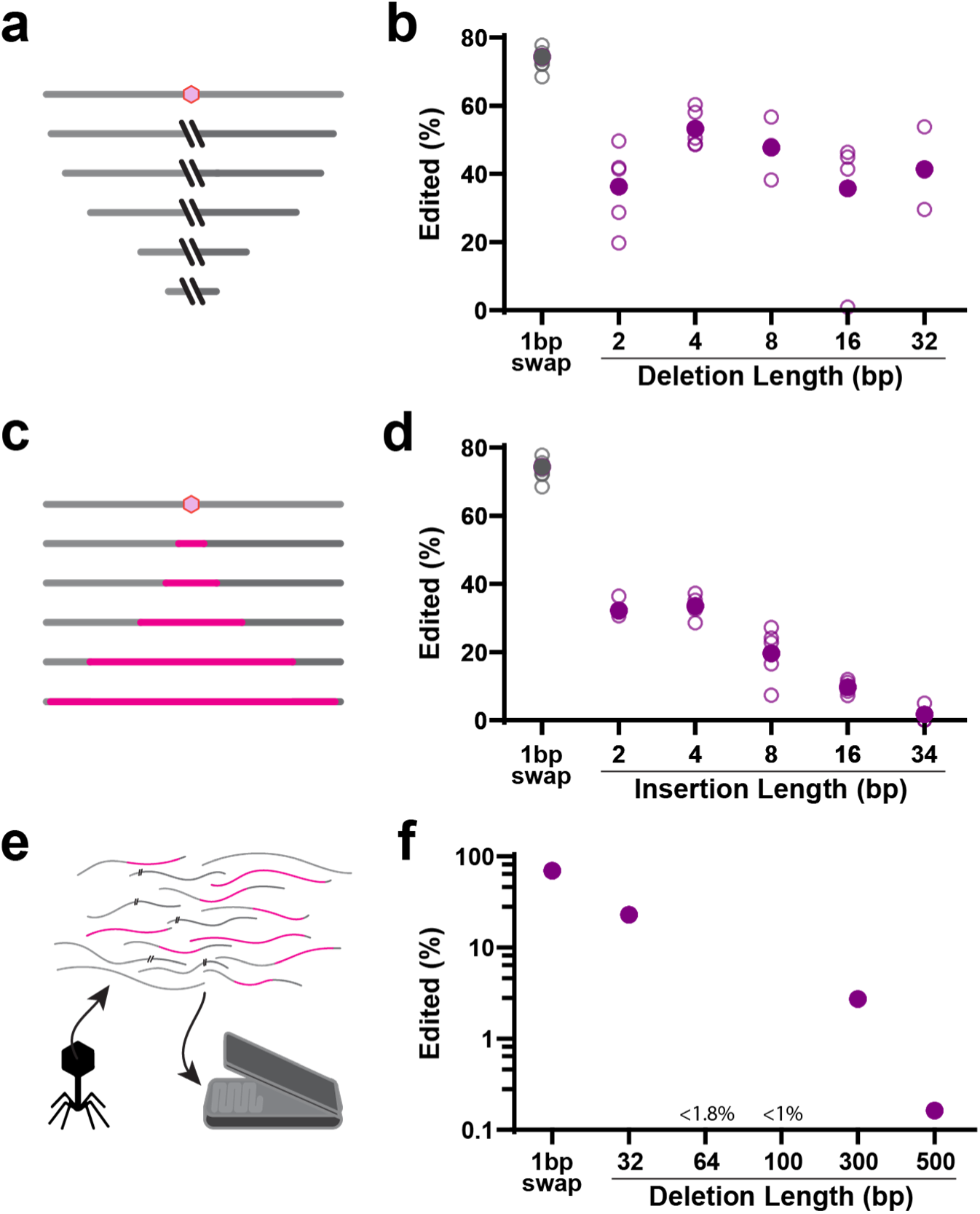
Insertions and Deletions via Recombitrons. **a.** Schematic of phage genome deletions of increasing size around a common central site (pink hexagon). **b.** Quantification of editing efficiency for deletions preceding position 37,673 as compared to a single base pair synonymous substitution. Five biological replicates are shown in open circles and the mean is shown as a closed circle. There is no significant effect of deletion length (one-way ANOVA, *P*=0.1564), but all deletions occur at a significantly lower frequency than the single-base substitution (one-way ANOVA, *P*<0.0001; Dunnett’s corrected vs 2 *P*<0.0001 vs 4 *P*=0.0083 vs 8 *P*=0.0008 vs 16 *P*<0.0001 vs 32 *P*=0.0005). **c.** Schematic of phage genome insertions of increasing size ending at a common position (pink hexagon). **d.** Quantification of editing efficiency for insertions at position 37,673 as compared to a single base pair synonymous substitution. Five biological replicates are shown in open circles and the mean is shown as a closed circle. Note that the same single base substitution was used as a parallel control for both deletions and insertions and thus the same points are shown here as in b. There is a significant effect of insertion length (one-way ANOVA, *P*<0.0001), and all insertions occur at a significantly lower frequency than the single-base substitution (one-way ANOVA, *P*<0.0002; Dunnett’s corrected *P*<0.0001 for all insertions). **e.** Schematic illustrating long-read nanopore quantification. **f.** Quantification of editing efficiency using nanopore sequencing. Points shown represent the fraction of edited reads pooled across three biological replicates. 64 and 100 base pair deletions were not detected in 57 and 102 reads aligned to the edit region, respectively. Additional statistical details in Supplementary Table 1.

We next tested insertion of 2, 4, 8, 16, or 32 bases into the same lambda site (**Fig 3c**). Here, we found a range of editing efficiencies, from ∼32% for insertions of 2 bases to ∼10% for insertions of 16 bases (**Fig 3d**). Like the deletions, all insertions were significantly less efficient than a synonymous single base substitution. Unlike the deletions, insertion size significantly affected efficiency, favoring smaller insertions.

Whereas the synonymous single base substitution is assumed to be neutral for phage fitness, we cannot assume the same for the deletions and insertions, which could affect transcription or phage packaging. This may underlie the lower, although still substantial, rate of deletions and insertions overall. Additionally, the PCR required for analysis of editing by amplicon sequencing is known to be biased toward smaller amplicons, which could inflate the measured rates of the larger deletions and decrease the measured rates of larger insertions.

To test yet larger insertions in a manner that is not subject to the size bias of PCR, we edited phages and then sequenced their genomes without amplification using long-read nanopore sequencing (**Fig 3e**). After editing, we isolated phage genomes via Norgen Phage DNA Isolation kit, attached barcodes and nanopore adapters, and sequenced molecules for 24-48h on a MinION (Oxford Nanopore Technologies). We quantified the resulting data using custom analysis software that binned reads by alignments to three possible genomes: a wild-type lambda genome, a lambda genome containing the relevant edit, or the BL21 *E. coli* genome as a negative control. BLAST alignment scores (percent identity, e-value, and alignment length) were compared for any read aligning to the lambda genome (wild-type or edited) in the region of the intended edit to quantify wild-type versus edited genomes. Consistent with a PCR bias for size, we found that amplification-free sequencing resulted in comparable editing rates to amplification-based sequencing for quantifying a single base substitution, but slightly lower rates of editing when measuring a 32 base pair deletion as compared to amplification-based sequencing (**Supplementary Figure 3a**).

This amplification-free sequencing approach enabled us to extend our deletion and insertion size ranges. For deletions, we tested 32, 64, 100, 300, or 500 base pairs. We found deletions of 32 bases at a frequency of ∼23%, 300 bases at a frequency of ∼2.7%, and 500 bases at a frequency of 0.16% (**Fig 3f**). We did not observe deletions of 64 or 100 bases in our data. However, our long-read coverage of the editing region in these samples was limited, so we can only conclude that, if these edits occur, they are present at <1.8% and <1% respectively across the three replicates of our sequencing data (**Supplementary Figure 3b,c**). In cases where we observe deletions, they were reliably found at the intended location and of the intended size (**Supplementary Figure 3e**).

We also tested insertion of larger sequences. We built recombitrons to insert a 34 base frt recombination site, a 264 base anti-CRISPR protein (AcrIIA4), a 393 base anti-CRISPR protein (AcrIIA13), and a 714 base sfGFP. As anticipated from the low insertion rates of insertions >8 base pairs, we did not observe any of these larger insertions in our long-read sequencing data, which was read-limited to a detection level of ∼1% (**Supplementary Figure 3d**). Thus, the recombitrons enable high efficiency deletions of up to 32 base pairs and insertions of up to 8 base pairs, but are not practical for counterselection-free isolation of larger deletions or insertions.

### Multiplexed Phage Engineering via Recombitrons

One shortcoming of all current phage editing approaches is an inability to simultaneously modify multiple sites in parallel on the same phage. Such parallel modification would be of great benefit to efforts aimed at engineering phages for targeted killing of pathogenic bacteria, but current approaches require cycles of editing, isolation, and re-editing that are impractical in an academic or industrial setting. We reasoned that a slight modification of our recombitron approach would enable such parallel, multiplexed editing of distant sites across a genome. In this modification, multiple bacterial strains that each harbor distinct recombitrons targeting different parts of the phage genome are mixed in a single culture. The phage to be edited is then propagated through that mixed culture. Every new infection event is an opportunity to acquire an edit from one of the recombitrons and, over time, these distinct, distant edits accumulate on individual phage genomes (**Fig 4a**). We ran these experiments in the engineered K-strain, through which the percentage of edited phage genomes increases with additional rounds of editing, beginning at a higher overall rate than the original experiments presented in Figure 1. Editing lambda in this strain yields efficiencies that reach >95% after three rounds of culture (**Fig 4b**).

**Figure 4.**
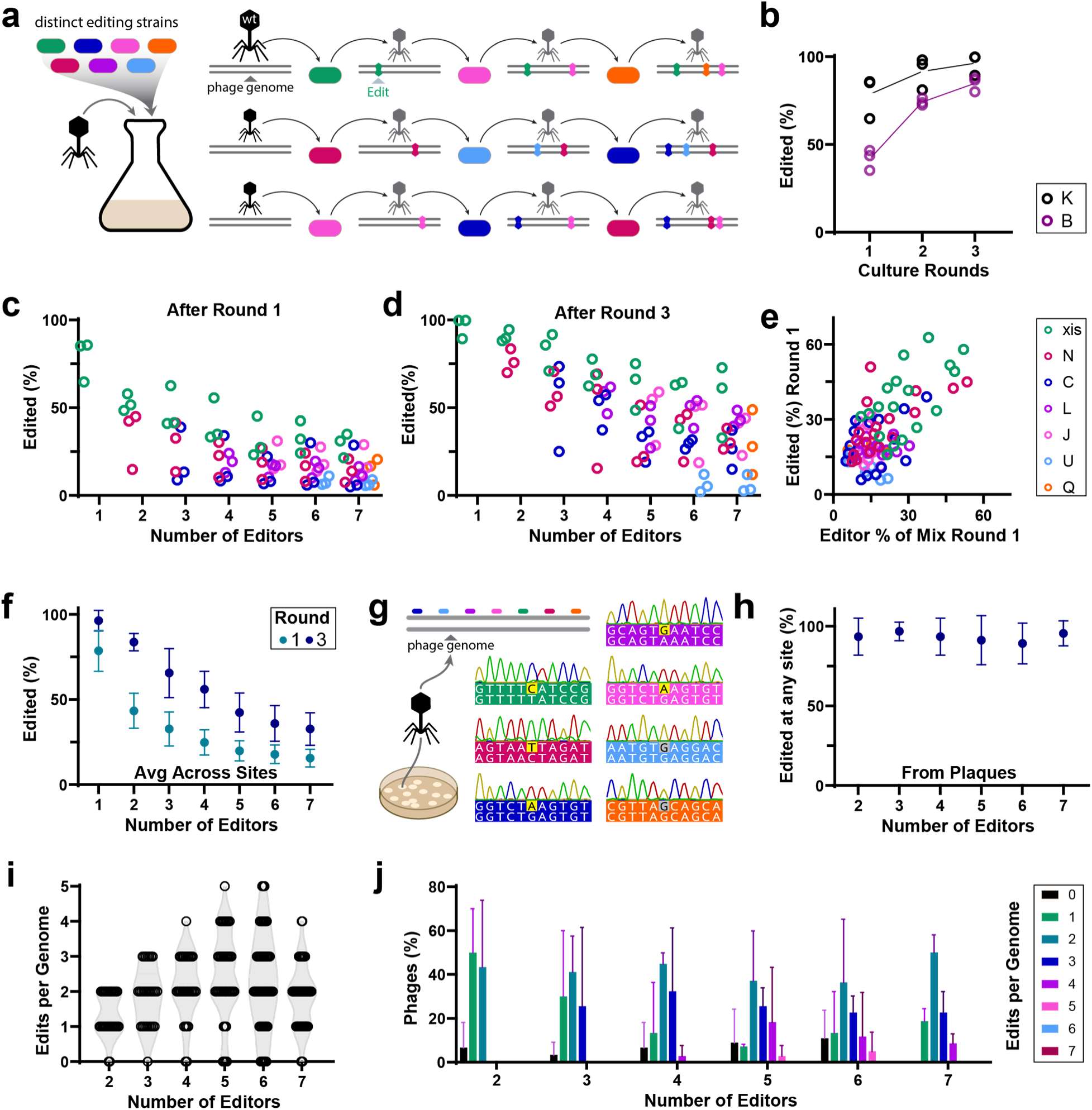
Multiplexed Phage Engineering via Recombitrons. **a.** Schematic illustrating the propagation of phages through cells expressing distinct recombitrons in a co-culture. Over generations of phage propagation, phage genomes accumulate multiple edits from different RT-Donors. **b.** Editing (%) increases over three rounds of editing. Editing in engineered K-strain *E. coli* outperforms the B-strain (two-way ANOVA, strain *P*<0.001, round *P*<0.0001, interaction *P*=0.0282). Three biological replicates are shown in open circles, lines connect averages. **c.** Editing (%) of each site from mixed recombitron cultures after 1 round of editing. Three biological replicates are shown in open circles for each site, clustered over the number of recombitrons used. **d.** Editing (%) of each site from mixed recombitron cultures after 3 rounds of editing, shown as in c. **e.** Editing (%) of each site after 1 round of editing in relation to the proportion of that recombitron strain in the culture (%) (two-tailed Pearson correlation, *P*<0.0001). **f.** Average editing (%) of all sites in each mixed culture. Points represent mean of three biological replicates ±SD (two-way ANOVA, number of editors *P*<0.0001, round *P*<0.0001, interaction *P*=0.234) **g.** Schematic illustrating Sanger sequencing of plaques, which enables quantification of each editing site from clonal phages. Representative data from one plaque with five edits (yellow) and two wild-type sites (grey). Panels **h**, **i**, and **j** are composed of data from three biological replicates. Total plaques: 2 editors, 30 plaques; 3 editors, 29 plaques; 4 editors, 31 plaques; 5 editors, 42 plaques; 6 editors, 53 plaques; 7 editors, 55 plaques **h.** Overall editing (%) of any site on plaques isolated from mixed cultures. Points represent mean of three biological replicates ±SD. **i.** Number of edits made on a single phage genome from mixed cultures, by number of recombitrons. Individual plaques from three biological replicates are shown as open circles on a violin plot of distribution. **j.** Number of edits made on a single phage genome from mixed cultures, by number of recombitrons, shown as a histogram (mean ±SD). Additional statistical details in Supplementary Table 1.

We created seven bacterial editing strains using the lambda recombitrons tested in Figure 1b. We then made mixed cultures of 1, 2, 3, 4, 5, 6, or all 7 bacterial strains and performed editing of lambda phage in these mixtures. These strains were grown individually in liquid culture overnight, then separately pre-induced for 2 hours, before being diluted to OD 0.25 and then mixed together in equal proportions. We infected these cultures with lambda (ΔcI) at an MOI of 0.1, grew the cultures for 16 hours overnight, and collected the phage lysate the next morning. We then used that phage stock to perform two additional rounds of editing in mixed-host cultures prepared identically to those of the first round. Based on our results that the amount of phage put into the culture is roughly the amount of phage extracted (**Supplementary Fig 1d,e)**, we added the same volume of phage in each round based on the titer determination before round one. We quantified editing across phage genomes and genomic loci for each of the mixtures using Illumina sequencing. We found editing across all sites from these mixed cultures after one round and increased editing in all cultures and sites after three rounds (**Fig 4c,d**). We found that the editing rate at any given site across all the mixtures was well correlated with the percentage of that editor in the mixture after round one (**Fig 4e**), and that the overall editing rate across all sites declined with the number of recombitron strains used (**Fig 4f**). This is consistent with a dilution effect of the strains on each other, which suggests that the overall editing rate is limited by the number of cells expressing each recombitron available for the phage to propagate through.

While amplicon sequencing showed substantial editing across many sites in a population of phages, we cannot use it to determine whether multiple edits are accumulating on individual phage genomes. Therefore, we plated phages after round three from each condition on bacterial lawns and Sanger sequenced individual plaques at each edit site (**Fig 4g**). This showed that 93.2% of phages were edited at one site or more, without employing any form of counter selection (**Fig 4h**). The plaque-based analysis strongly mirrored the Illumina sequencing analysis of editing rate per site by mixture, when plaque data for a condition was pooled (**Supplementary Fig 4**). With the plaque data, however, we can also look at edits per phage genome. We found that multiple sites were edited on a majority of the phages across all conditions, with individual phages edited at up to 5 distinct locations (**Fig 4i,j**). This represents a new milestone in the continuous, multiplexed, selection-free engineering of phage genomes.

## DISCUSSION

Here we present a new approach to phage editing using recombitrons that leverages modified retrons to continuously provide RT-Donors for recombineering in phage hosts. Recombitrons enable counter-selection-free generation of phage mutants across multiple phages, with optimized forms yielding up to 100% editing efficiency. Moreover, recombitrons can be multiplexed to generate multiple distant mutations on individual phage genomes. Critically, this approach is easy to perform. Recombitrons are generated from simple, standard cloning methods via inexpensive, short oligonucleotides. The process of editing merely requires propagating the bacteria/phage culture, with no intervening transformations or special reagents. Once recombitrons are cloned and phage stocks are prepared, the generation of lambda phage edited at up to five distinct positions required hands-on time of less than 2 hours.

This technical advance is poised to change the way we approach innovation in phage biology and phage therapeutics. For instance, studying epistasis in phage genomes becomes a feasible experiment that merely requires mixing recombitron strains in different combinations. The technical hurdle of phage library generation is also dramatically reduced. In our multiplexed experiment, 93.2% of phages were edited. Host range is a major determinant of phage therapy efficacy, which could be addressed by rapidly screening a large library targeting the tail fiber or other critical host range genes for a fraction of the effort of current approaches.

Some outstanding questions and further engineering challenges remain. For instance, overexpression of a compatible SSB provided a large benefit to the editing of T7, which demonstrates that recombitrons can be optimized for generality across phages. However, that effect was not immediately transferable to T5. Presumably, T5 may be limited by another phage-specific requirement, and T2 is presumably limited by the use of modified nucleotides. Further work will seek to engineer around these idiosyncrasies and increase the extensibility of this approach to other phages. Finally, in future work we will explore the use of this technology beyond *E. coli* phages.

## METHODS

Biological replicates were taken from distinct samples, not the same sample measured repeatedly.

### Bacterial Strains and Growth Conditions

This work used the following *E. coli* strains: NEB 5-alpha (NEB, C2987; not authenticated), BL21-AI (Thermo Fisher, C607003; not authenticated), bMS.346 and bSLS.114. bMS.346 (used previously^20^) was generated from *E. coli* MG1655 by inactivating the *exoI* and *recJ* genes with early stop codons. bSLS.114 (used previously^29^) was generated from BL21-AI by deleting the retron Eco1 locus by lambda Red recombinase-mediated insertion of an FRT-flanked chloramphenicol resistance cassette, which was subsequently excised using FLP recombinase^42^. bCF.5 was generated from bSLS.114, also using the lambda Red system. A 12.1kb region was deleted that contains a partial lambda*B prophage that is native to BL21-AI cells within the attB site, where temperate lambda integrates into the bacterial genome^43^.

Phage retron recombineering cultures were grown in LB, shaking at 37 °C with appropriate inducers and antibiotics. Inducers and antibiotics were used at the following working concentrations: 2 mg/ml L-arabinose (GoldBio, A-300), 1 mM IPTG (GoldBio, I2481C), 1mM m-toluic acid (Sigma-Aldrich, 202-723-9), 35 µg/ml kanamycin (GoldBio, K-120), 100 µg/ml carbenicillin (GoldBio, C-103) and 25 µg/ml chloramphenicol (GoldBio, C-105; used at 10 µg/ml for selection during bacterial recombineering for strain generation).

### Plasmid Construction

All cloning steps were performed in *E. coli* NEB 5-alpha. pORTMAGE-Ec1 was generated previously (Addgene plasmid no. 138474)^27^. Derivatives of pORTMAGE-Ec1 (pCF.109, pCF.110, pCF.111) were cloned to contain an additional SSB protein, amplified with PCR from its host genome, via Gibson Assembly. Plasmids for RT-Donor production, containing the retron-Eco1 RT and ncRNA with extended a1/a2 regions, were cloned from pSLS.492. pSLS.492 was generated previously (Addgene plasmid no. 184957)^20^. Specific donor sequences for small edits were encoded in primers and substituted into the RT-DNA-encoding region of the ncRNA with a PCR and KLD reaction (NEB M0554). Donor sequences for larger insertions were cloned through Gibson assembly, using synthesized gene fragments (Twist Biosciences). Recombitron ncRNAs encoding the editing donors are listed in **Supplementary Table 2**.

### Phage Strains and Propagation

Phages were propagated from ATCC stocks (Lambda #97538, Lambda WT #23724-B2, T7 #BAA-1025-B2, T5 #11303-B5, T2 #11303-B2) into a 2mL culture of *E. coli* (BL21*^ΔEco^*^1^) at 37°C at OD600 0.25 in LB medium supplemented with 0.1 mM MnCl_2_ and 5 mM MgCl_2_ (MMB) until culture collapse, according to established techniques^44,45^. The culture was then centrifuged for 10 min at 4000 rpm and the supernatant was filtered through a 0.2μm filter to remove bacterial remnants. Lysate titer was determined using the full plate plaque assay method as described by Kropinski et al.^46^. We used recombitrons to edit this lambda strain to encode two early stop codons in the cI gene, responsible for lysogeny control, to ensure the phage was strictly lytic (lambda ΔcI). After recombineering, we Sanger sequenced plaques to check the edit sites. We isolated an edited plaque and used Illumina Miseq of its lysate to ensure purity of the edited phage. We used this strictly lytic version for all experiments involving lambda phage, unless otherwise noted.

Genomic locations used to label edits are from wild-type reference sequences of phages available through NCBI GenBank: lambda (J02459.1), T5 (AY587007.1), T7 (V01146.1), and T2 (AP018813.1). We found that the strain of phage lambda we used naturally contains a large genomic deletion between 21738 and 27723. This region encodes genes that are not well-characterized, but may be involved in lysogeny control^47,48^.

### Plaque Assays

Small drop and full plate plaque assays were performed as previously described by Mazzocco et al.^49^, starting from bacteria grown overnight at 37°C. For small drop plaque assays, 200ul of the bacterial culture was mixed with 2mL melted MMB agar (LB + 0.1 mM MnCl_2_ + 5 mM MgCl_2_ + 0.75% agar) and plated on MMB agar plates. 10-fold serial dilutions in MMB were performed for each of the phages and 2ul drops were placed on the bacterial layer. The plates were dried for 20 min at room temperature and then incubated overnight at 37°C. Full plate plaque assays were set up by mixing 200ul of the bacterial culture with 20ul of phage lysate, using 10-fold serial dilutions of the lysate to achieve between 200-10 plaques. After incubating at room temperature without shaking for 5 min, the mixture was added to 2mL melted MMB agar and poured onto MMB agar plates. The plates were dried for 20 min at room temperature and incubated overnight at 37°C. Plaque forming units were counted to calculate the titer.

### Recombineering and Sequencing

The retron cassette, with modified ncRNA to contain a donor, was co-expressed with CspRecT and mutL E32K from the plasmid pORTMAGE-Ec1. All experiments, except multiplexing and large insertions/deletions (>30 bp), were conducted in 500uL cultures in a deep 96-well plate. Multiplexing experiments were conducted in 3mL cultures in 15mL tubes. For amplification-free sequencing by nanopore, larger culture volumes (25mL cultures in 250 mL flasks) were used to enable collection of a higher quantity of phage DNA. Cultures were induced for 2 hrs at 37 °C, with shaking. The OD600 of each culture was measured to approximate cell density and cultures were diluted to OD600 0.25. Phages were originally propagated through the corresponding host that would be used for editing (B- or K-strain E coli). A volume of pre-titered phage was added to the culture to reach a multiplicity of infection (MOI) of 0.1. The infected culture was grown overnight for 16 hrs, before being centrifuged for 10 min at 4000 rpm to remove the cells. The supernatant was filtered through a 0.2μm filter to isolate phage.

For amplicon-based sequencing, the lysate was mixed 1:1 with DNase/RNase-free water and the mixture was incubated at 95°C for 5 min. This boiled culture (0.25ul) was used as a template in a 25ul PCR reaction with primers flanking the edit site on the phage genome. These amplicons were indexed and sequenced on an Illumina MiSeq instrument. Sequencing primers are listed in **Supplementary Table 3**.

For amplification-free sequencing, extracellular DNA was removed through DNase I treatment, with 20 U of DNase I (NEB, M0303S) for 1mL of phage lysate, incubated at room temperature for 15 min and then inactivated at 75°C for 5 min. Phage were then lysed and DNA extracted using the Norgen Phage DNA Isolation Kit (Norgen, 46800). The samples were prepped for sequencing in a standard Nanopore workflow. DNA ends were repaired using NEBNext Ultra II End Repair/dA-Tailing Module (NEB, E7526S). End-repaired DNA was then cleaned up using Ampure XP beads. Barcodes were ligated using the standard protocol for Nanopore Barcode Expansion Kit (Oxford Nanopore Technologies, EXP-NBD196). After barcoding, the standard Oxford Nanopore adaptor ligation, clean-up, and loading protocols were followed for Ligation Sequencing Kit 109 and Flow Cell 106 for the MinION instrument (Oxford Nanopore Technologies, SQK-LSK109, FLO-MIN106D). Base calling was performed using Guppy Basecaller with high accuracy and barcode trimming settings.

Sanger sequencing of phage plaques was accomplished by picking plaques produced from the full plate assay described above. Plates were sent to Azenta/Genewiz for sequencing with one of the MiSeq-compatible primers used to assess the same site. Sequences were analyzed using Geneious through alignment to the region surrounding the edit site on the phage genome.

### Editing Rate Quantification

A custom Python workflow was used to quantify edits from amplicon sequencing data. Reads were required to contain outside flanking nucleotide sequences that occur on the phage genome, but beyond the RT-Donor region to avoid quantifying RT-DNA or plasmid. Reads were then trimmed by left and right sequences immediately flanking the edit site. Reads containing these inside flanking sequences in the correct order with an appropriate distance between them (depending on edit type) were assigned to either wild-type, edit, or other. The edit percentage is the number of edited reads over the sum of all reads containing flanking sequences.

A distinct custom Python program was used for quantifying amplification-free nanopore sequencing data, due to the higher error rates in nanopore sequencing and the lack of a defined region of the genome contained in each read. Reads were aligned using BLAST+ to three reference genomes: wild-type lambda, edited lambda containing the matching edit to the read’s experiment, and BL21-AI *E. coli*. If reads aligned to either lambda genomes, the read’s alignment coordinates had to be at least 50 bases past the insertion/deletion coordinates, as well as be > 500 bp and have >50% of the read mapped to the reference genome as quality scores. If a read aligned to the insertion/deletion point and passed all quality scores, the percent identity and alignment length over read length were compared to assign the read as either wild-type or edited. Coverage of the edit region was 50-1000x per experimental condition.

### Data Availability

All data supporting the findings of this study are available within the article and its supplementary information, or will be made available from the authors upon request. Sequencing data associated with this study are available in the NCBI SRA (PRJNA933262).

### Code Availability

Custom code to process or analyze data from this study will be made available on GitHub prior to peer-reviewed publication.

## Acknowledgements

Work was supported by funding from the National Science Foundation (MCB 2137692), the National Institute of Biomedical Imaging and Bioengineering (R21EB031393), the Gary and Eileen Morgenthaler Fund, and the National Institute of General Medical Sciences (1DP2GM140917). S.L.S. acknowledges additional funding support from the L.K. Whittier Foundation and the Pew Biomedical Scholars Program.

## Author Contributions

C.B.F., S.B.-K., and S.L.S. conceived the study and, with K.D.C., K.A.Z., and A.G.-D., outlined the scope of the project and designed experiments. C.B.F. developed phage handling and editing protocols. Experiments were performed and analyzed by C.B.F. (Fig1 e-h, Fig2, Fig4, Supp Fig1 b-h, Supp Fig2, Supp Fig4), K.D.C. (Fig3, Supp Fig3), S.B.-K. (Fig1 b-d, Supp Fig1 a), and K.A.Z (Fig3, Supp Fig3). C.B.F. and S.L.S. wrote the manuscript, with input from all authors.

## CORRESPONDING AUTHOR

Correspondence to Seth L. Shipman.

## Supplementary Figures

**Supplementary Figure 1.**
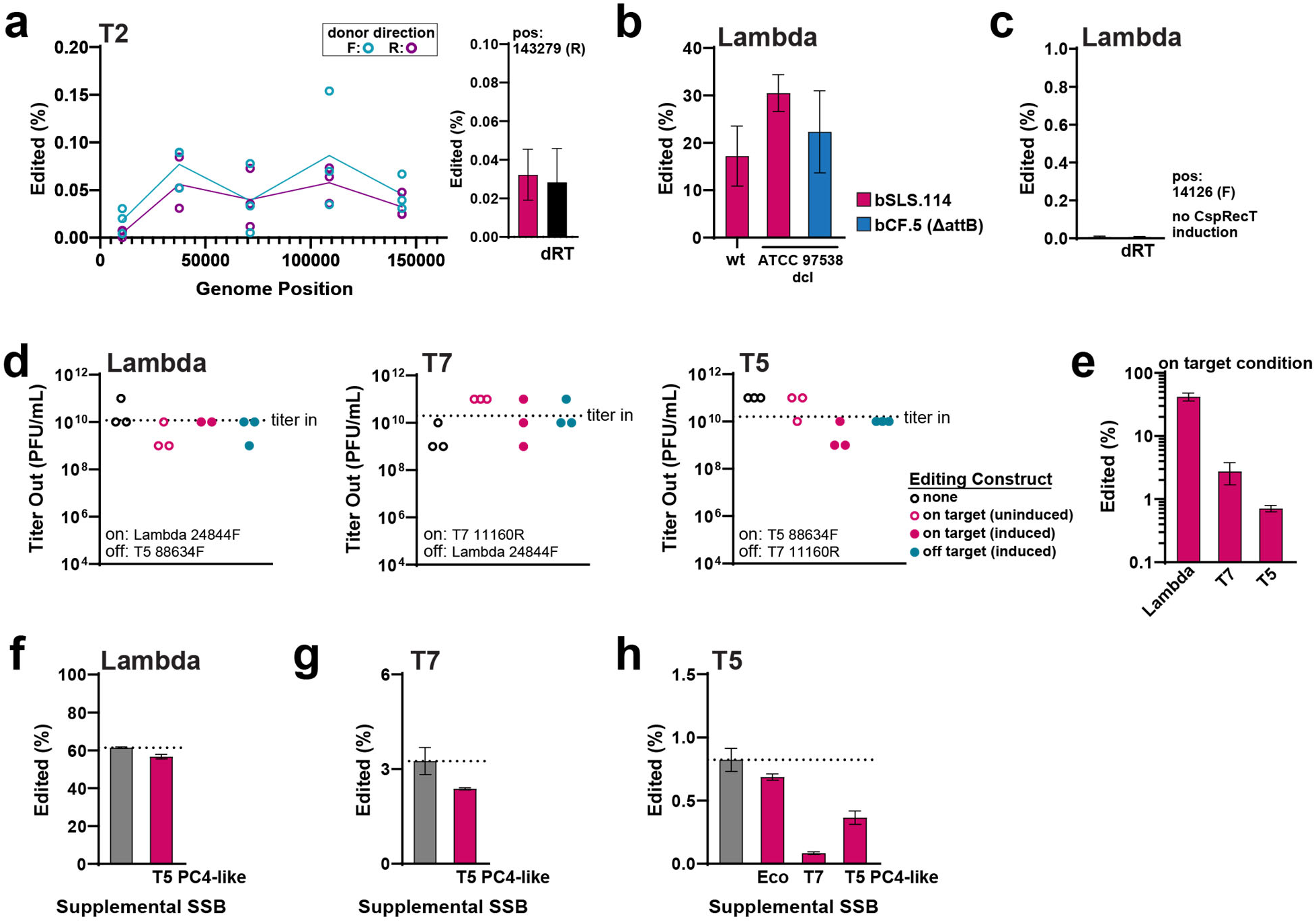
Accompaniment to Figure 1. **a.** Left: Edited phage genomes (as a % of all genomes) across phage T2. Editing with a forward RT-DNA is shown in blue, reverse in purple. Three individual replicates are shown in open circles at each point. Right: Editing with the recombitron at site 143,279 (R) (±SD) versus a dRT control. **b.** Editing (%) of lambda and lambda_ΔcI in different host strains at site 14,070 (R). Mean ±SD is shown for 3 (wt in bSLS.114), 5 (ΔcI in bSLS.114), and 2 (ΔcI in bCF.5) biological replicates. **c.** Editing (%) of lambda without induction of CspRecT. Mean ±SD for 3 biological replicates at site 14,126. **d.** Titer (PFU/mL) of phage lambda, T7, and T5 after propagation through host cells of different conditions, compared to amount of phage added to the culture. Host conditions are without recombitrons (open black circles), with uninduced recombitrons (open pink circles), with induced recombitrons (closed pink circles), and with induced recombitrons that target a different phage (closed blue circles). Individual biological replicates are shown in circles for each condition. **e.** Editing (%) of phage lambda, T7, and T5 from the induced, on-target recombitron condition in panel d. **f.** Editing (%) of lambda at site 14126 (F) compared to editing with supplemental expression of T5 SSB. **g.** Editing (%) of T7 at site 22872 (R) compared to editing with supplemental expression of T5 SSB. **h.** Editing (%) of T5 at site 88634 (F) with supplemental expression of *E. coli* SSB, T7 SSB, or T5 SSB.

**Supplementary Figure 2.**
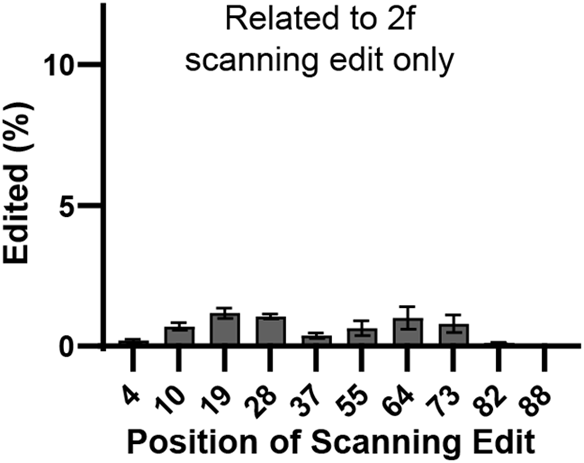
Accompaniment to Figure 2. Rate of acquiring only the scanning edit in lambda when donors contain both scanning and central edits. (mean ±SD).

**Supplementary Figure 3.**
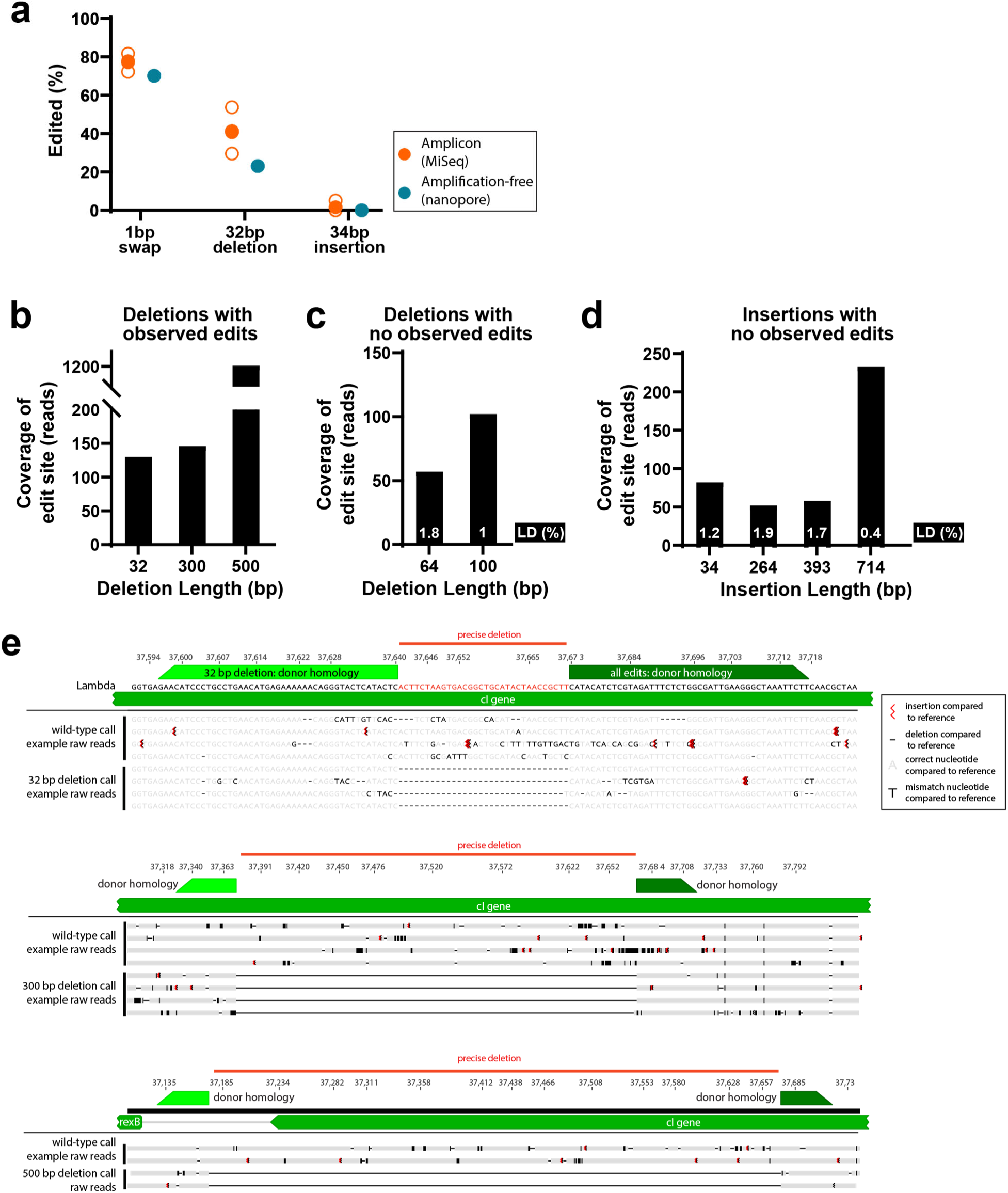
Accompaniment to Figure 3. **a.** Comparison of edited phages measure by amplicon (Illumina) or amplification-free (Oxford Nanopore) sequencing. Open orange circles represent biological replicates of amplicon data and filled orange circle represents the mean. Filled blue circle represents the aggregate nanopore data from three replicates. **b.** Coverage of the editing site in long-read nanopore sequencing for deletions in which we observe editing. **c.** Coverage of the editing site in long-read nanopore sequencing for deletions in which we do not observe any edits. Estimated limit of detection for these samples is calculated by dividing 100 by the coverage of the site. **d.** Coverage of the editing site in long-read nanopore sequencing for large insertions, for which we do not observe any edits. Estimated limit of detection for these samples is calculated by dividing 100 by the coverage of the site. **e.** Examples of nanopore reads for different deletion conditions.

**Supplementary Figure 4.**
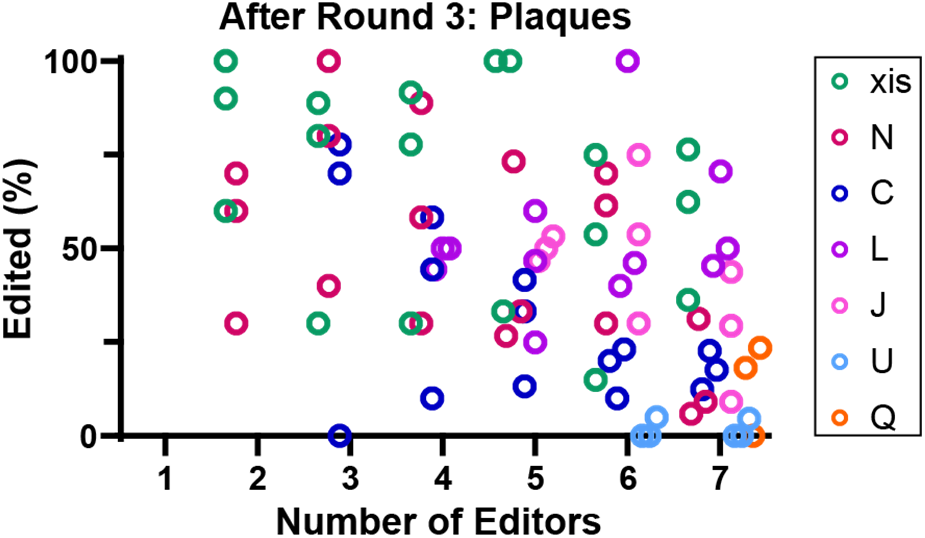
Accompaniment to Figure 4. Editing (%) from Sanger sequencing of plaques at each site from mixed recombitron cultures after 3 rounds of editing. Three biological replicates are shown in open circles for each site, clustered over the number of recombitrons used.

